# Recombinant human soluble thrombomodulin is associated with attenuation of sepsis-induced renal impairment by inhibition of extracellular histone release

**DOI:** 10.1101/780403

**Authors:** Masayuki Akatsuka, Yoshiki Masuda, Hiroomi Tatsumi, Michiaki Yamakage

## Abstract

Multiple organ dysfunction induced by sepsis often involves kidney injury. Extracellular histones released in response to damage-associated molecular patterns are known to facilitate sepsis-induced organ dysfunction. Recombinant human soluble thrombomodulin (rhTM) and its lectin-like domain (D1) exert anti-inflammatory effects and neutralize damage-associated molecular patterns. However, the effects of rhTM and D1 on extracellular histone H3 levels and kidney injury remain poorly understood. Our purpose was to investigate the association between extracellular histone H3 levels and kidney injury, and to clarify the effects of rhTM and D1 on extracellular histone H3 levels, kidney injury, and survival in sepsis-induced rats. Rats in whom sepsis was induced via cecal ligation and puncture were used in this study. Histone H3 levels, histopathology of the kidneys, and the survival rate of rats at 24 h after cecal ligation and puncture were investigated. Histone H3 levels increased over time following cecal ligation and puncture. Histopathological analyses indicated that the distribution of degeneration foci among tubular epithelial cells of the kidney and levels of histone H3 increased simultaneously. Administration of rhTM and D1 significantly reduced histone H3 levels compared with that in the vehicle-treated group and improved kidney injury. The survival rates of rats in rhTM- and D1-treated groups were significantly higher than that in the vehicle-treated group. The results of this study indicated that rhTM and its D1 similarly reduce elevated histone H3 levels, thereby reducing acute kidney injury. Our findings also proposed that rhTM and D1 show potential as new treatment strategies for sepsis combined with acute kidney injury.

## Introduction

Sepsis is a life-threatening healthcare issue caused by infection, and associated with a high risk of mortality. In sepsis, inflammatory response occurs following infection and sometimes contributes to the development of various organ dysfunctions. Kidney injury is one of the most common organ dysfunctions induced by sepsis. Once a sepsis patient develops complications of acute kidney injury (AKI), mortality is reportedly increased [1-3].

In recent years, extracellular histones have been attracting attention as mediators of death from sepsis [4]. Extracellular histones are known as damage-associated molecular patterns (DAMPs) and are released when cellular injury occurs, resulting in induction of neutrophil migration, platelet aggregation, and endothelial cell damage [5, 6]. DAMPs are likely to trigger the activation of serious inflammation [7] and result in severe tissue injury [8-12]. Suppression of the production of inflammatory mediators such as cytokines and DAMPs could thus contribute to the improvement of clinical outcomes in critically ill patients.

Thrombomodulin (TM) is a thrombin-binding anticoagulant cofactor that is expressed on the surface of endothelial cells and plays an important role in the regulation of intravascular coagulation [13]. The structure of TM consists of five domains: the N-terminal lectin-like domain (D1), a domain with six epidermal growth factor (EGF)-like structures (D2), a serine and threonine-rich domain (D3), a transmembrane domain (D4), and a short cytoplasmic domain (D5) [14]. Recent studies have shown that D1 has anti-inflammatory effects, inhibits the production of inflammatory cytokines, and also binds to high-mobility group box-1 (HMGB1), which is a known damage-associated molecular pattern (DAMP) [15, 16]. However, whether administration of TM could suppress extracellular histone H3 levels and improve organ injury remains unclear. In addition, whether administration of D1 has the same effects on histone H3 levels and protective effects against kidney injury remains to be clarified. We hypothesized that administration of TM and D1 would improve kidney injury and survival rate in sepsis-induced rats by inhibiting extracellular histone H3 release. Recombinant human soluble TM (rhTM) is homologous with the extracellular domains of TM and is used as an anticoagulant in treatment of disseminated intravascular coagulation in Japan. The purposes of this study were: 1) to investigate the relationship between extracellular histone H3 levels and kidney injury; and 2) to clarify whether rhTM and D1 could affect extracellular histone H3 levels, kidney injury, and survival rate of sepsis-induced rats.

## Materials and methods

### Cecal ligation and puncture (CLP)-induced sepsis model in rats

Male Wistar rats were obtained from Japan SLC, Inc. (Hamamatsu, Shizuoka, Japan). The rats weighing 250-300 g were used after an acclimation period of at least 1 week before experimentation. Rats were maintained in a 12/12 h light/dark cycle to minimize noises, vibrations and odors. The animals were kept in temperatures of 22-24°C with 40-50% humidity. Food and clean water were freely accessible at all times. Therefore, not all animal welfare considerations taken were necessary, including efforts to minimize suffering and distress, use of analgesics or anesthetics, or special housing conditions. To create an in vivo sepsis-induced model, we performed CLP procedures as follows. Rats were anesthetized with isoflurane, and a 1.5-cm incision was made along the abdominal midline. Ligation was performed at a distance of 1-2 cm from the blind-ending cecum, avoiding the mesenteric vessels of the cecum. The distal cecum was then punctured once with an 18-G needle, and a small amount of feces was gently squeezed out. The cecum was returned to the abdominal cavity, and layers of the abdominal incision were sequentially sutured. The sham rats were anesthetized and underwent the same procedure without CLP. After the surgical procedure, rats were subcutaneously injected with 10 mL/kg of saline. Rats were deprived of food and free access to water after CLP.

rhTM and D1 were used for experimental studies at the indicated dose, which was provided by Asahi Kasei Pharma (Tokyo, Japan). The molecular weights of rhTM and D1 are 52,171 Da and 20,191 Da, respectively. The dose of D1 in this study was thus calculated as 0.36 of the rhTM dose by conversion according to relative molecular weights for equivalent titration.

Blood samples were collected by cardiac puncture in EDTA-containing blood collection tubes at each time point under general anesthesia and centrifuged at 1500×*g* for 10 min. The supernatant after centrifugation was collected and stored at -80°C until further analyses.

All experimental protocols described in the present study were approved by the Institutional Animal Care and Use Committee of Sapporo Medical University (Permit No. 19-023). The staff who conducted this experiment had taken a special seminar and training in animal care or handling that were approved by the ethics committee of our university. All efforts were made to minimize animal suffering. In the present study, as the humane end point for rats, a state of respiratory urgency, continuous laying, or convulsion was judged as indicative of a moribund state and treated with euthanasia. Once an animal reached endpoint criteria, they were euthanized immediately and counted as an animal death. Euthanasia was performed by deep anesthesia with isoflurane, causing irreversible cardiac or respiratory arrest. Animal health and behavior were observed carefully every 30 minutes after the CLP procedure. Additionally, animal welfare was checked every 4 h until 24 h after the procedure.

In the first line of experiments, we examined the levels of circulating histone H3 and effective concentrations of rhTM. The sample size was determined to be at least 5 rats per group via power analysis based on a previous study [17]. We used new rats every 4 h from 0 to 24 h, with 5 rats in each group: sham, vehicle, low-, middle-, and high-dose groups (Figs 1 and 2). In the second line of experiments, we used 6 animals in each group to verify the effects of both rhTM and D1 on histone H3 levels after CLP (Figs 3-7). In the third line of experiments, we used 10 animals in each group to confirm the pharmacological effects of these proteins (rhTM and D1) on animal models to check animal welfare (Fig 8). A total of 124 rats were used in this study and all of them were euthanized after meeting the endpoint criteria within 24 h. Data from all rats that underwent experimentation were included in this study, with none excluded.

**Fig 1.**
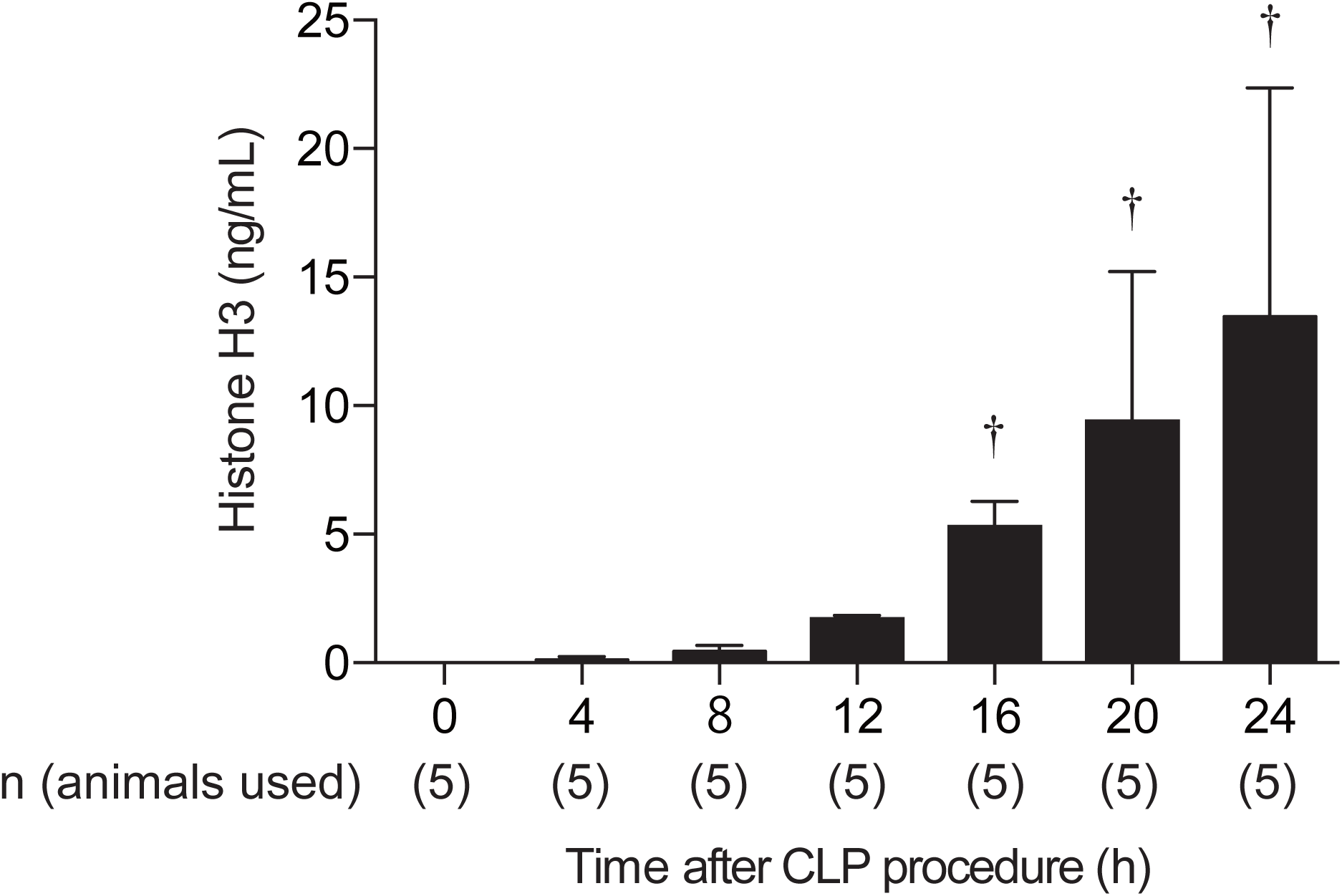
Circulating histone H3 level after CLP procedure. Circulating histone H3 level elevated over time. n = 5 per group. † *P* ≤ 0.01 vs. 0 h.

**Fig 2.**
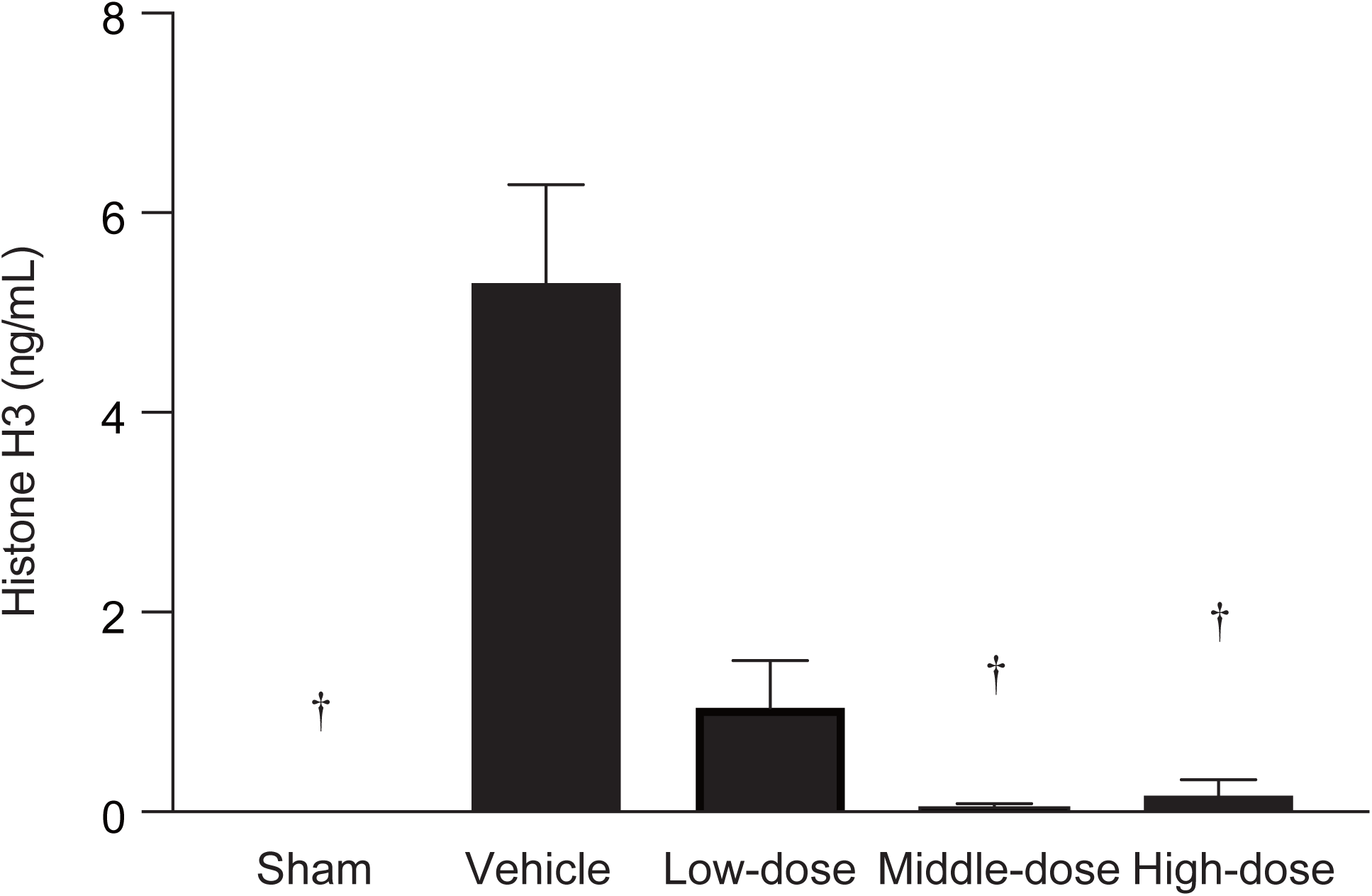
Effectiveness of rhTM for reducing histone levels. Circulating histone H3 levels were significantly lower with administration of middle- or high-dose (3 mg/kg or 6 mg/kg) rhTM compared to control group administered vehicle. n = 5 per group. †*P* ≤ 0.01 vs. vehicle.

**Fig 3.**
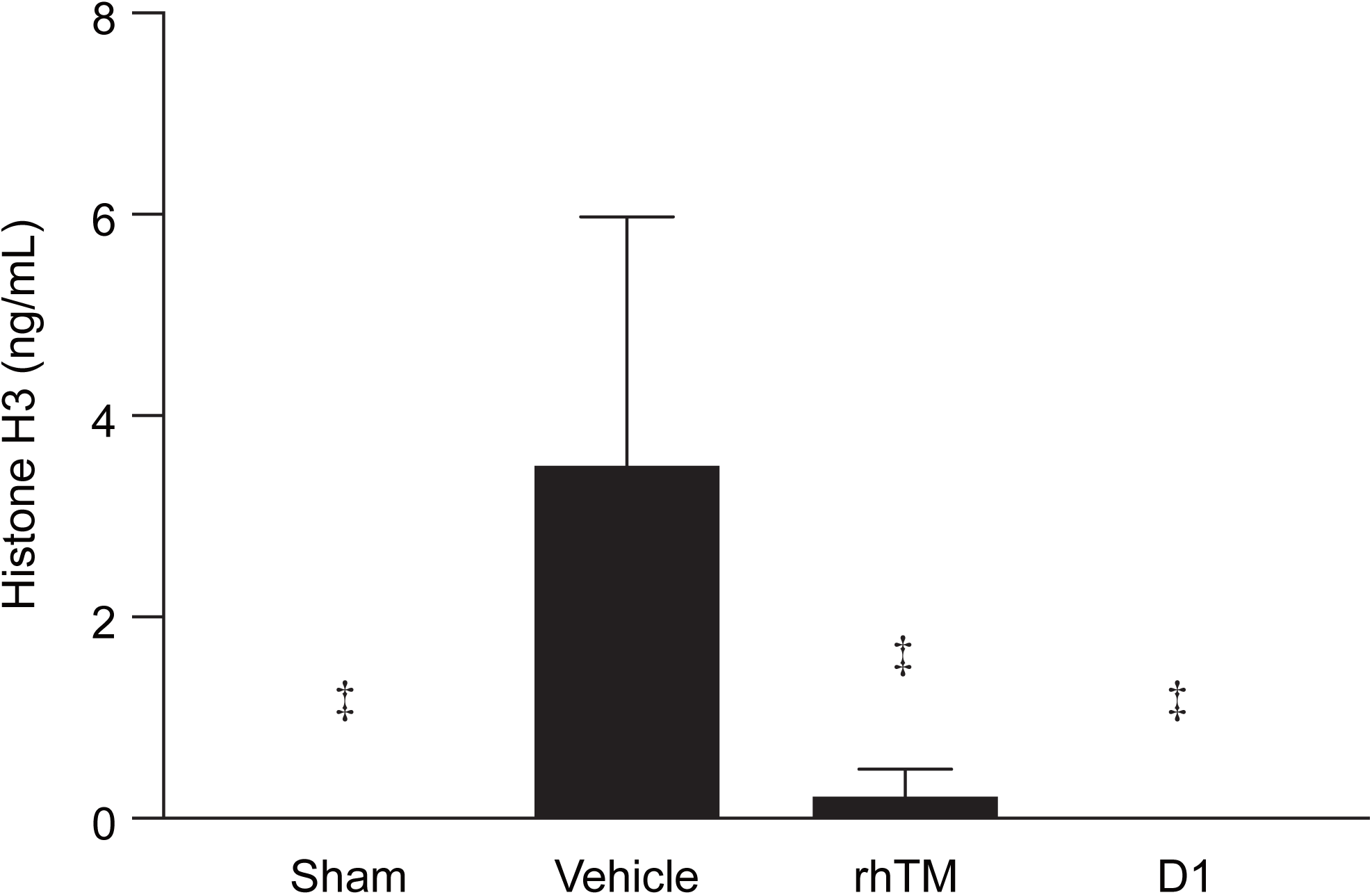
Effect of lectin-like domain 1 (D1) of rhTM on reduction of histone H3 levels in sepsis-induced rats. Administration of rhTM and D1 significantly decreased circulating histone H3 levels. n = 6 per group. ‡ *P* ≤ 0.001 vs. vehicle.

**Fig 4.**
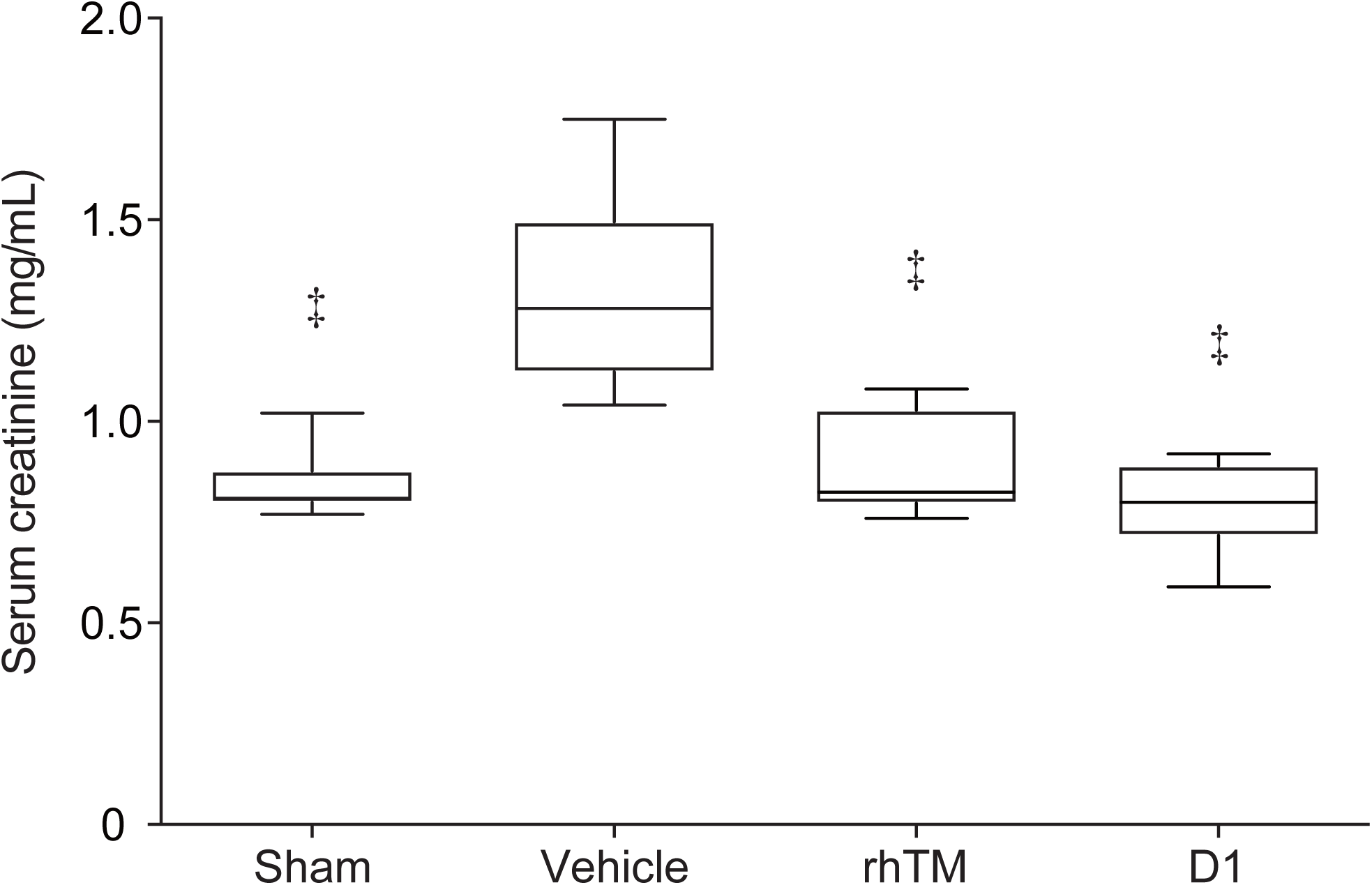
Serum creatinine level in rats after CLP procedure. Administration of rhTM and D1 improved serum creatinine levels. n = 6 per group. ‡ *P* ≤ 0.001 vs. vehicle.

**Fig 5.**
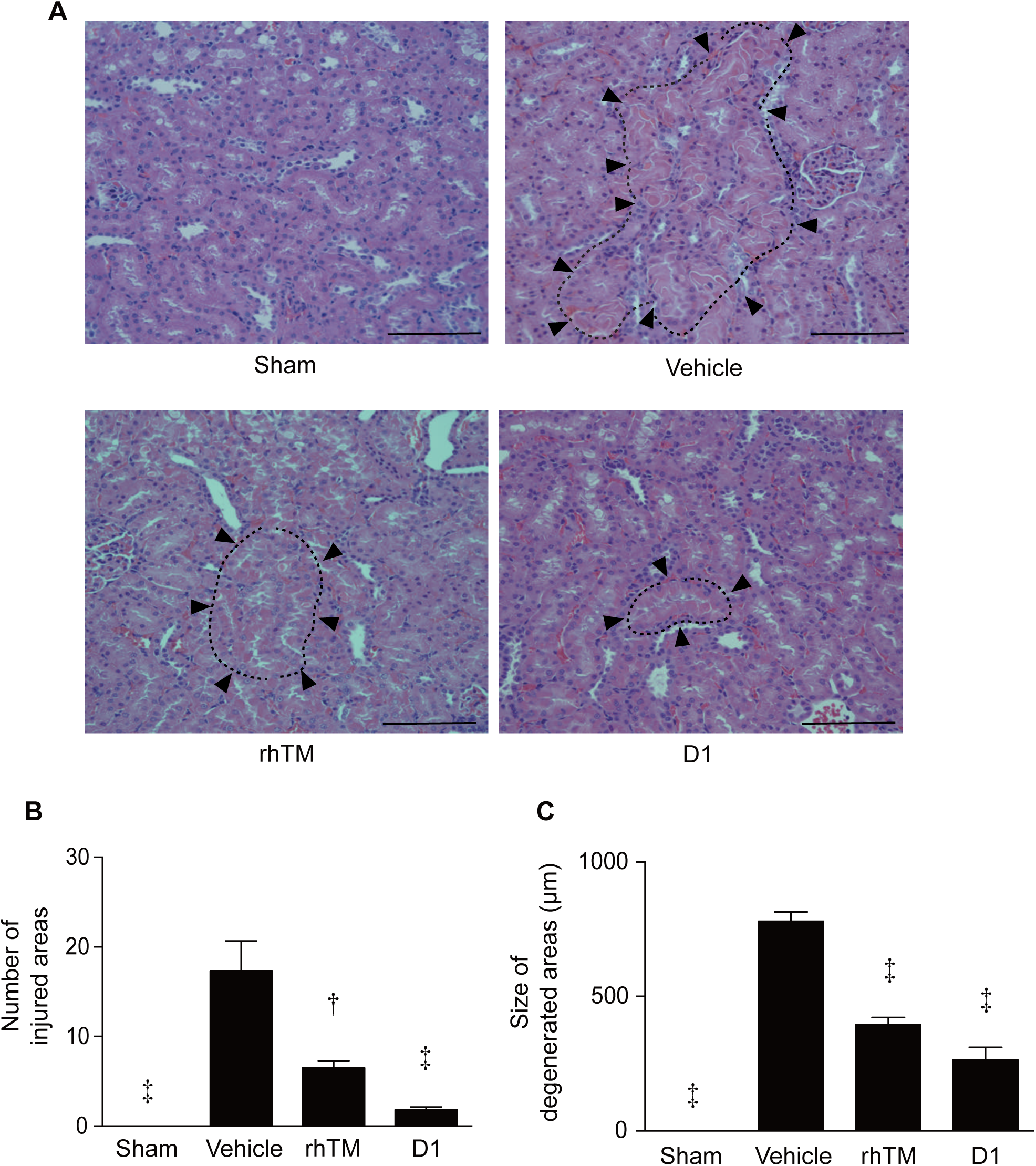
Histopathologic examination of kidney stained with HE. A) Histologic findings in rat kidney. Renal tissue was obtained 16 h after CLP and stained with HE. Arrowheads indicate degenerated areas. Representative images are shown at a magnification of ×200. B) Quantification of kidney degeneration in the four groups is shown (number of injured areas). N = 6 per group. C) Quantification of kidney degeneration in the four groups (size of degenerated areas). n = 6 per group. † *P* ≤ 0.01 vs. vehicle. ‡ *P* ≤ 0.001 vs. vehicle. Scale bars = 200 µm.

**Fig 6.**
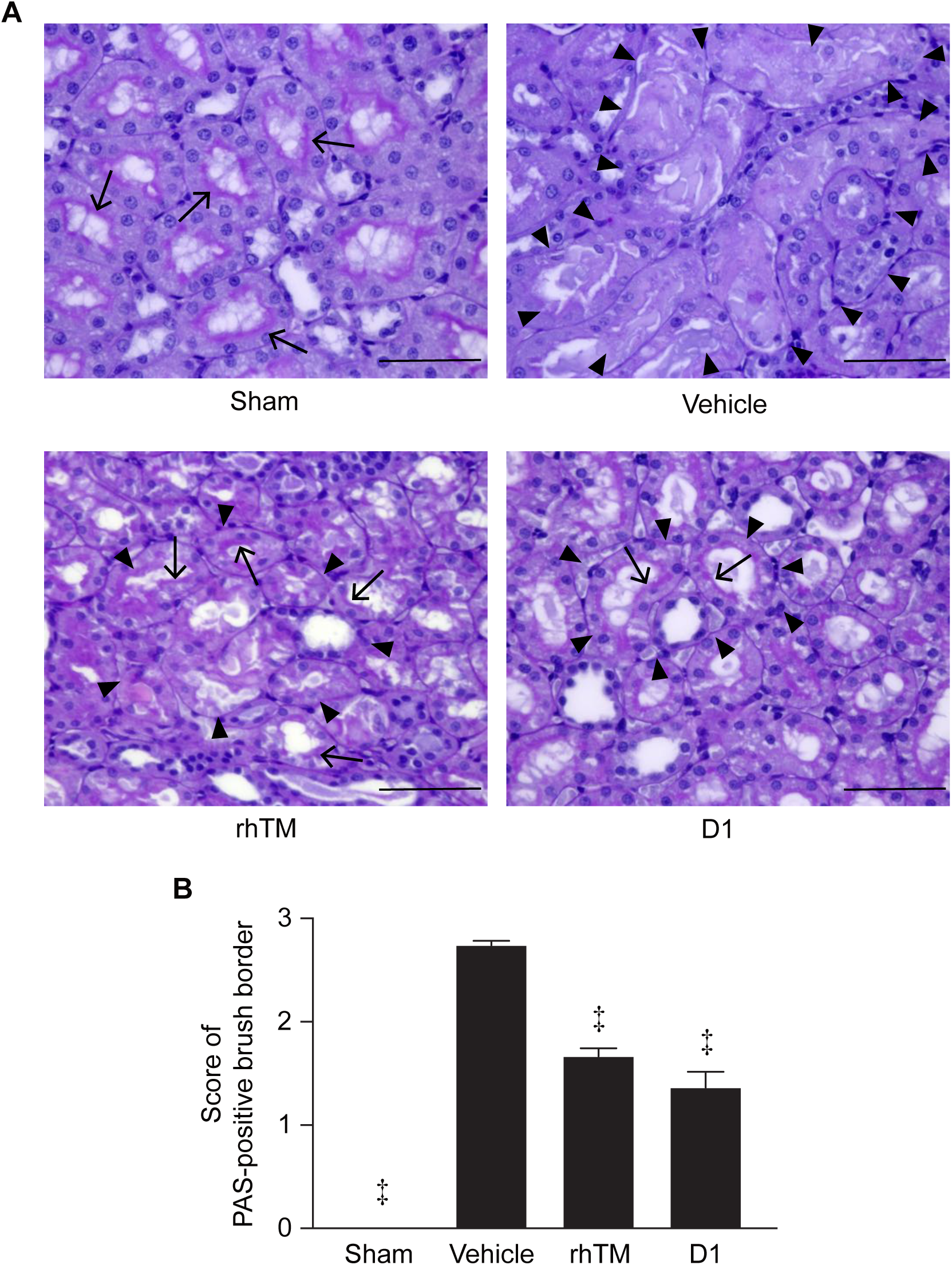
Histopathologiccal examination of kidney stained with PAS. A) Histologic findings in rat kidney. Renal tissue was obtained 16 h after CLP and stained with PAS. Arrowheads indicate degenerated areas and arrows indicate the PAS-positive brush border. Representative images are shown at a magnification of ×400. B) The score of the PAS-positive brush border in the four groups is shown (score of PAS=positive brush border). n = 6 per group. ‡ *P* ≤ 0.001 vs. vehicle. Scale bars = 100 µm.

**Fig 7.**
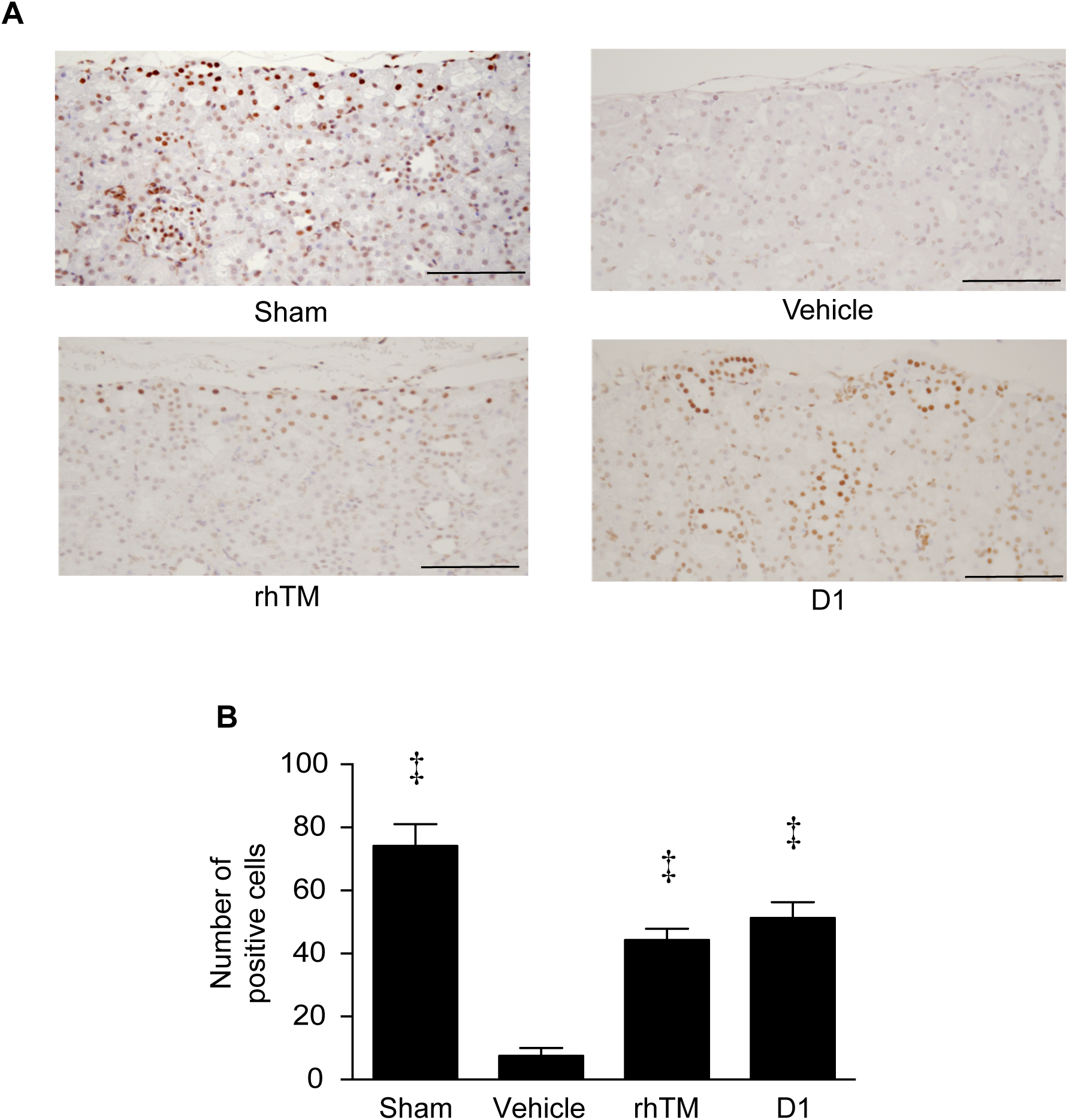
Histopathologic examination of kidney with immunohistochemical staining for histone H3. A) Immunohistochemical findings in rat kidney. Renal tissue was obtained 16 h after CLP and stained with antihistone H3 peptide polyclonal antibody. Representative images are shown at a magnification of ×200. B) Quantification of positive cells for immunohistochemical staining in the four groups is shown. n = 6 per group. ‡ *P* ≤ 0.001 vs. vehicle. Scale bars = 200 µm.

**Fig 8.**
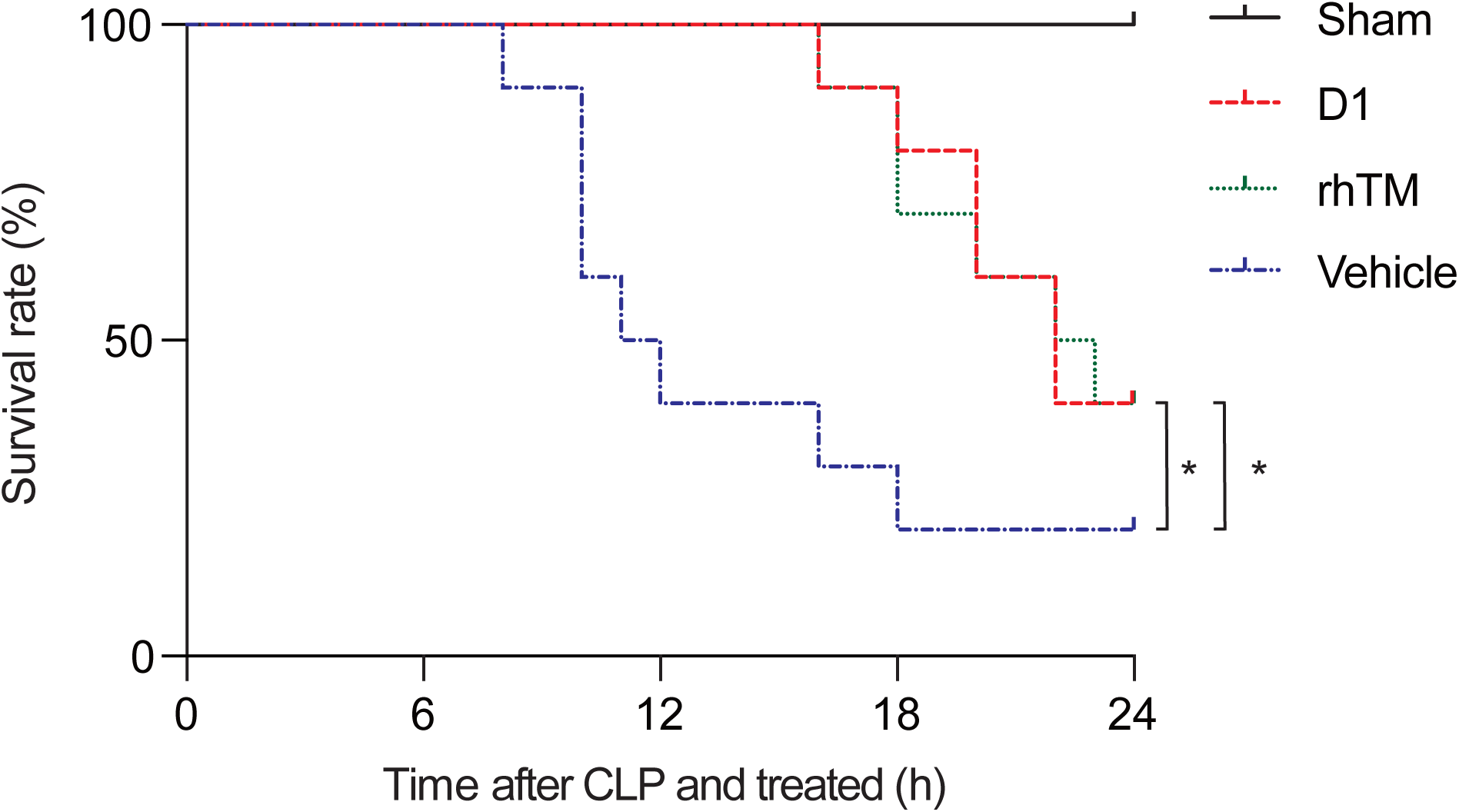
Survival curves in rats after CLP procedure. Rats were administered either: rhTM (green dotted line); D1 (red dashed line); or saline as a vehicle (blue dashed dotted line). Sham rats are represented by the black solid line. n = 10 per group. * *P* < 0.05 vs. vehicle.

### Determination of histone H3 levels after CLP

We investigated changes in circulating histone H3 levels after CLP. Blood samples were collected from rats under general anesthesia. We observed the rats every 4 h until 24 h after the end of the CLP procedure. Thus, the duration of the experiment was set at 24 h. We determined the timing of elevations in histone H3 levels. Commercial enzyme-linked immunosorbent assay kits were used for measurement of histone H3 levels (Shino-Test, Kanazawa, Japan) according to the instructions from the manufacturer.

### Administration of rhTM after CLP or optimal dose of rhTM for affecting histone H3 levels after CLP

We administered several doses of rhTM after CLP as a bolus injection into the tail vein to determine the optimal dose of rhTM for affecting histone H3 levels. rhTM was administered through the tail vein immediately after the CLP procedure. We set four groups: a control group that received 2 mL of 0.9% saline as a vehicle; and low-, middle-, and high-dose groups that received rhTM at 1 mg/kg, 3 mg/kg, and 6 mg/kg, respectively. Blood samples were obtained at the time point of significant increases in histone H3 levels after the CLP procedures in each group and histone H3 levels were compared between these four groups.

### Administration of D1 after CLP

To evaluate the efficacy of D1 on histone H3 levels, 0.9% saline as a vehicle and a calculated dose of D1 based on the optimal dose of rhTM were administered as a bolus injection into the tail vein of rats after CLP. Blood samples were collected from rats under general anesthesia at the time point of a significant increase in histone H3 levels after CLP. Histone H3 levels were then determined and compared between vehicle- and D1-treated groups.

### Determination of serum creatinine level

We evaluated serum creatinine level using blood samples obtained from the following groups: sham, vehicle, rhTM, and D1. Serum creatinine concentrations in plasma were measured using detection kits (Arbor Assays, Cat. #KB02-H1) according to the manufacturer’s instructions.

### Histopathologic analysis of kidney

Kidney tissue specimens were obtained after euthanasia by deep anesthesia and were submerged in 10% formaldehyde neutral buffer solution. Fixed tissues were embedded in paraffin, cut at intervals of 3 µm, and stained with hematoxylin and eosin (HE). The number and size of injured areas in the whole kidney were determined.

Formalin-fixed hemisected kidneys were embedded in paraffin and stained with periodic acid-Schiff (PAS). Histological analysis of kidney tubular cell damage was performed using the following scoring assessment: 0, no damage; 1, mild (PAS-positive brush border that is lost in <1/3 of the injured area); 2, moderate (<2/3); 3, severe (complete loss or ≥2/3).

### Immunohistochemical staining of the kidney and quantification

Immunohistochemical staining for histone H3 was performed using rabbit polyclonal antibody (Lot. 001; Shino-Test). After antigen retrieval and blocking, sections were preincubated with 10% normal goat serum for 20 min. Sections were incubated with a 1:500 dilution of the primary antibody histone H3 for 15 min. Sections were then incubated with Bond™ Polymer Refine Detection (Leica Biosystems, Newcastle-Upon-Tyne, UK). The chromogen was 3,3’-diaminobenzidine (DAB), followed by counterstaining with hematoxylin. Staining intensity was graded as: 0, negative; 1, weak; 2, moderate; or 3, strong. A grade of 2 or 3 was considered positive for immunohistochemical staining, whereas a grade of 0 or 1 was considered negative. We counted the positive cells in a randomly selected microscopic field at high magnification (×400). Quantitative analysis of histone H3 expression was performed using histological images that were digitally obtained using a light microscope in order to best reflect overall immunostaining of the specimen on each slide. Camera control with constant setup conditions was maintained throughout the course of the study. Adobe Photoshop software (Adobe Systems, Mountain View, CA) was used to process acquired images. For quantification, positive signals were selected using a constant threshold in each image at high magnification (×400). All immunohistochemical specimens were evaluated by a board-certified pathologist (Department of Pathology, Sapporo Medical University School of Medicine, Japan).

### Survival analysis

To evaluate the effects of rhTM and D1 on survival rate after CLP, rats were divided into four groups: a sham group of rats without CLP; a control group of rats administered vehicle after CLP; an rhTM group of rats administered the optimal dose of rhTM after CLP; and a D1 group of rats administered D1 (0.36 of rhTM dose) after CLP. Survival of the rats was monitored up to for 24 h. We analyzed and compared survival rates of rats between these four groups.

### Statistical analysis

Data are presented as the mean ± standard error of the mean (SEM). Statistical analysis between groups was performed by Tukey’s multiple comparison test. Survival rate was analyzed by the Kaplan-Meier method using log-rank testing. Analyses were performed with Graph Pad Prism version 6.0h software (GraphPad Software, San Diego, CA). The level of significance was set at 0.05 for all analyses using two-tailed *P* values.

## Results

### Serial changes in extracellular histone H3 levels after CLP procedure

Circulating histone H3 level began to significantly increase at 16 h after CLP and continued to increase up to 24 h (Fig 1). Based on this result for this timing, we decided to evaluate extracellular histone H3 levels at 16 h after CLP in the following studies.

### Effect of rhTM on histone H3 levels after CLP

Significant differences were found in circulating histone H3 levels at 16 h after CLP among the four groups. Histone H3 levels in middle-dose (3 mg/kg) and high-dose (6 mg/kg) rhTM groups were significantly lower than that in the control group administered vehicle only (Fig 2). Therefore, in the following experiment, we used rhTM dose at 3 mg/kg for evaluation of rhTM efficacy on decreasing histone H3 level, improving survival rate and protective actions on the kidney.

### Effect of D1 on histone H3 levels after CLP

The administration dose of D1 was set at 1.09 mg/kg converted according to the molecular weight of rhTM. Administration of D1 significantly reduced circulating histone H3 levels, similar to rhTM (Fig 3). No significant difference in reduction of histone H3 levels was seen between rhTM-treated and D1-treated groups.

### Effect of rhTM and D1 on serum creatinine levels

Serum creatinine levels were significantly lower in rhTM- and D1-treated rats than in vehicle-treated rats (Fig 4). No significant difference in serum creatinine level was seen between rhTM and D1 groups.

### rhTM and D1 improve histopathologic findings of the kidney

After CLP in rats, scattered degeneration foci were widely observed in tubular epithelial cells and were widely distributed throughout the whole kidney in rats in the vehicle-treated group (Figs 5A-C). We observed that more severe degenerations were mainly observed in the tubular epithelium, suggesting peripheral circulatory insufficiency (Fig 5A). On the other hand, in rats treated with rhTM and D1, tubular structure was maintained and degeneration was observed in small areas. As a result of quantification, administration of rhTM and D1 significantly decreased numbers and sizes of areas of degeneration in the kidney compared with non-treated rats (Figs 5B and 5C). Also, we observed almost no or a small portion of the brush border with PAS staining in the tubular lumen of kidneys in the vehicle group (Fig 6A). In contrast, the structure of the brush border in the rhTM and D1 groups was maintained in large areas, although there was partial injury to the tubular lumen (Fig 6A). The histological scoring for the degree of kidney tubular damage in the rhTM and D1 groups was significantly lower than the vehicle group (Fig 6B).

In rats from the sham group, immunostaining for histone H3 was mainly observed in the cortical area. However, in rats from the vehicle-treated group, less immunostaining for histone H3 was observed compared with the sham group (Fig 7A). Rats administered and D1 showed more immunostaining for histone H3 compared with the vehicle-treated group. We performed quantitative analysis of histone H3 expression in the cortical area for each group (Fig 7B). The number of positive cells in the non-treated group was significantly decreased compared with the sham group. Administration of rhTM and D1 significantly increased the number of positive cells compared with the vehicle-treated group. Our data suggested that soluble TM can rescue sepsis-induced renal tubular damage.

### Survival rate

The survival rate at 24 h after CLP was 40% in the rhTM-treated group and 20% in the vehicle-treated group. The difference in survival rate between the two groups was significant according to Kaplan-Meier analysis (*P* = 0.048) (Fig 8). In the D1-treated group, the survival rate for 24 h after the CLP procedure was 40%, compared with 20% in the vehicle-treated group. Kaplan-Meier analysis revealed significant prolongation of survival in the D1-treated group compared with the vehicle-treated group (*P* = 0.038) (Fig 8).

## Discussion

The present study showed two important findings. First, circulating histone H3 levels elevated over time and were associated with kidney injury diagnosed pathologically and from renal function, and with high mortality in rats with intraabdominal infection-induced sepsis. Second, we found that administration of rhTM and D1 dramatically reduced circulating histone H3 levels, protected against kidney injury, and improved mortality in comparison with non-treated rats with intraabdominal sepsis. From the present results, administration of rhTM appears to be associated with renal protective effects via attenuation of histone H3 levels and consequent improvement of survival rate in sepsis rats. The lectin-like domain of rhTM, which does not show anticoagulant activity, is suggested to play a central role in anti-inflammatory activity. rhTM, and the D1 domain in particular, thus holds promise as a drug for the treatment and prevention of septic AKI.

In this study, rhTM and D1 were administered at the time of CLP production. We found that suppression of the increase in histone H3 levels is associated with improvement of kidney injury. This shows the possibility that AKI is associated with increased serum histone H3 levels. In the pathological examination of renal tissue, AKI was induced by CLP and a significant decrease in intracellular histone protein levels was identified. However, whether the decrease in intracellular histone H3 levels following AKI and the increase in serum histone H3 levels are related remains unclear. In addition, it is not clear how circulating histone H3 causes regional inflammation and cytotoxicity in the kidney. However, in general, leukocytes activated by inflammation are known to release histone H3, resulting in the elevation of histone H3 levels in sepsis [18]. Moreover, histones, which are released into the extracellular space through the process of cell death, cause kidney injury by direct cytotoxicity to renal tubular epithelium and direct interaction with toll-like receptor (TLR)-2 (TLR2) and TLR4 [19]. Another study using the ischemia-reperfusion model in mice showed that extracellular histones lead to multiple organ injury including kidney injury [20]. These histone-induced kidney injuries are consistent with our data. In this study, there were no injuries to organs other than the kidney in histological findings in rats with sepsis by CLP. In addition, histone H3 levels were increased in sepsis-induced rats. From these two observations, we surmised that increased histone H3 levels might be related to kidney injury. As for inflammatory markers, we measured tumor necrosis factor-α (TNF-α) after the CLP procedure. TNF-α levels increased earlier than the point when circulating histone H3 level increased, and the peaked out (S1 Fig). Therefore, TNF-a did not increase at the time when circulating histone H3 increased and kidney injury occurred. In terms of kidney injury markers, we measured serum creatinine levels as they are used as diagnostic criteria for acute kidney injury (Fig 4).

Conversely, Ito et al. [21] reported that depletion of leukocytes by anticancer drug-induced bone marrow suppression resulted in inhibition of histone H3 production in a mouse model of sepsis. Therefore, the increase in histone H3 levels in sepsis is thus highly possible to be derived from leukocytes. Since this study showed pathological findings that CLP induced disappearance of intracellular histone H3 in degenerated renal cells, damaged renal cells were considered to release intracellular histone H3 into the extracellular space and this mechanism is partially associated with increased serum histone H3 levels in sepsis rats.

Nuclear histone proteins have been reported to be released into the extracellular space and act as major mediators of death in sepsis [4, 22]. In our sepsis model, extracellular histone H3 levels increased significantly from 16 h after CLP. At that time, renal tissue showed extensive degeneration and vacuolation with increased histone H3 levels. However, no findings of acute tubular necrosis were identified. These findings show the early stage of AKI in pathological evaluations.

Results of histological findings demonstrated microcirculatory disturbance, showing the ischemic injury of kidney. Moreover, mortality rates of CLP-induced sepsis rats in the non-treated group, in which histone H3 levels were high, were significantly higher than in the rhTM and D1 treated groups. These results suggest that elevated histone H3 levels might be associated with kidney injury and prognosis in intraabdominal sepsis-induced rats.

rhTM has been reported to bind to DAMPs, resulting in reduction of DAMPs levels through inactivation by neutralization or decomposition. This elimination of released histone H3 has been demonstrated to improve mortality in experimental investigations [23-25]. rhTM comprises the following three structural regions: the lectin-like domain, the EGF-like domain, and a serine-threonine-rich region. The lectin-like domain of the three structures has an anti-inflammatory effect and a binding effect on HMGB1, a known DAMP [26]. Another study showed that rhTM (particularly the lectin-like domain) protects the kidney from ischemia-reperfusion injury via activated protein C (APC)-dependent and APC-independent mechanisms [27]. On the other hand, endothelial dysfunction during ischemic injury caused by inflammatory response results in microvascular congestion, leading to microcirculation abnormalities [28, 29]. Endothelial cells play an important role in preventing intravascular thrombus formation and maintain microcirculation [30]. However, the inhibitory effect of rhTM and D1 on circulating histone H3 level remains poorly understood. We therefore hypothesized that regulation of extracellular histone H3 by administration of rhTM and D1 would prevent sepsis-induced endothelial dysfunction, improve disturbances in microcirculation, and result in improved renal injury and prognosis in a rat sepsis model.

In our experimental study, CLP induced a significant increase in circulating extracellular histone H3 levels and induced AKI as evaluated pathologically and from serum creatinine levels. Results of immunohistochemical staining of rat renal cells also demonstrated decreased expression of histone H3. These data suggest that CLP-induced histone H3 elevation is associated with induction of AKI in rats.

Our results demonstrated that administration of rhTM and D1 inhibited CLP-induced histone H3 elevation and mitigated CLP-induced AKI. Moreover, administration of both rhTM and D1 improved prognosis of CLP-induced rats. Furthermore, results of immunohistochemical staining clearly demonstrated increased expression of histone H3 in rats that received either rhTM or D1, preventing histone H3 from being released to the extracellular space. These results suggest that rhTM and D1 might have the main effect of inhibiting the release of histone H3. On the other hand, no significant differences were evident between the rhTM-treated group and the D1-treated group in terms of the effects on inhibiting histone H3, improving renal injury, and prognosis. The main domain of rhTM that exerts anti-inflammatory action thus appears likely to be the lectin-like domain, D1.

The present study clearly demonstrated an association between increased histone H3 levels and kidney injury, and indicated that administration of rhTM and D1 reduced histone H3 levels and protected renal tissue. However, the following issues remain to be clarified in future studies: 1) the mechanisms underlying suppression of histone H3 by rhTM and D1; and 2) effects of rhTM and D1 on other organ injuries that were not examined in this study. Further studies are thus needed in the future to clarify the mechanisms regulating histone H3 and protecting against organ injuries by rhTM and D1.

In the future, further studies are required to clarify the relationship between disappearance of intracellular histone protein in renal tubular cells and increases in blood concentrations.

## Conclusions

CLP induced increases in extracellular histone levels, with this elevation peaking in the late phase of sepsis followed by the peak in tumor necrosis factor (TNF)-α level. CLP also induced AKI and this elevation of extracellular histone is thought to be related to CLP-induced AKI. rhTM and its lectin-like domain D1 were found to similarly reduce these elevated histone H3 levels, thereby protecting against CLP-induced AKI and significantly improving the survival rate of sepsis rats. Based on these facts, rhTM is considered to have anti-inflammatory activity through the suppression of histone H3 levels, and thus may have renal protective activity. Furthermore, since the D1 domain, which has no anticoagulant action among the domains of rhTM, has the same effects on reducing histone H3 levels as rhTM did, these findings suggest that the D1 domain may play a crucial role in anti-inflammatory activity among the domains of rhTM. The results of our study suggest that rhTM and D1 domains may be useful drugs for treating sepsis complicated by AKI. Our results also may provide a rationale for further clinical studies of rhTM and its lectin-like domain as a potential drug for the treatment of septic AKI.

## Acknowledgments

We wish to thank Professor Makoto Osanai for the histopathologic evaluations.

## Author Contributions

**Conceptualization:** Masayuki Akatsuka, Yoshiki Masuda.

**Data curation:** Masayuki Akatsuka, Yoshiki Masuda.

**Formal analysis:** Masayuki Akatsuka, Hiroomi Tatsumi.

**Funding acquisition:** Masayuki Akatsuka.

**Investigation:** Masayuki Akatsuka, Yoshiki Masuda.

**Methodology:** Masayuki Akatsuka, Yoshiki Masuda.

**Project administration:** Yoshiki Masuda, Michiaki Yamakage.

**Resources:** Masayuki Akatsuka, Hiroomi Tatsumi.

**Software:** Masayuki Akatsuka, Yoshiki Masuda.

**Supervision:** Yoshiki Masuda, Michiaki Yamakage.

**Validation:** Masayuki Akatsuka, Hiroomi Tatsumi, Yoshiki Masuda.

**Visualization:** Masayuki Akatsuka, Hiroomi Tatsumi.

**Writing – original draft:** Masayuki Akatsuka.

**Writing – review & editing:** Masayuki Akatsuka, Yoshiki Masuda, Michiaki Yamakage.

