## Supplementary material for "Recombinant human soluble thrombomodulin is associated with attenuation of sepsis-induced renal impairment by inhibition of extracellular histone release": S1 Fig

### Supporting information

**S1 Text. Determination of TNF-α levels after CLP.** We investigated changes in TNF-α levels after CLP. TNF-α concentrations was measured using the rat TNF-α quantikine ELISA kit (R&D Systems, MN, USA) according to the manufacturer's instructions.

**S1 Fig. TNF-α level after CLP procedure.** TNF-α level increased up to 8 h after the CLP procedure and then decreased over time. † $P \leq 0.01$ vs. 0 h. ‡ $P \leq 0.001$ vs. 0 h.

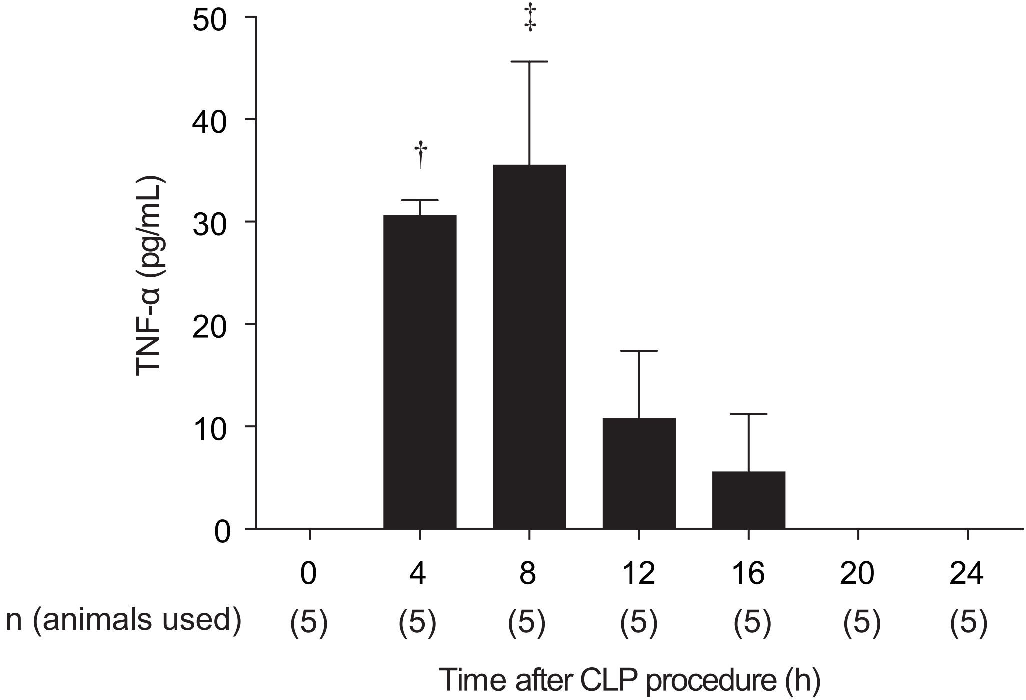
50
40
30
20
10
0
TNF-α (pg/mL)
†
‡
0
4
8
12
16
20
24
n (animals used)
(5)
(5)
(5)
(5)
(5)
(5)
(5)
Time after CLP procedure (h)

S1 Fig
